# Shear Stress Induces a Time-Dependent Inflammatory Response in Human Monocyte-Derived Macrophages

**DOI:** 10.1101/2022.12.08.519590

**Authors:** Elysa Jui, Griffin Kingsley, Hong Kim T. Phan, Kavya L. Singampalli, Ravi K. Birla, Jennifer P. Connell, Sundeep G. Keswani, K. Jane Grande-Allen

## Abstract

Macrophages are innate immune cells that are known for their extreme plasticity, enabling diverse phenotypes that lie on a continuum. In a simplified model, they switch between pro-inflammatory (M1) and anti-inflammatory (M2) phenotypes depending on surrounding microenvironmental cues, which have been implicated in disease outcomes. Although considerable research has been focused on macrophage response to biochemical cues and mechanical signals, there is a scarcity of knowledge surrounding their behavior in response to shear stress. In this study, we applied varying magnitudes of shear stress on human monocyte-derived macrophages (MDMs) using a cone-and-plate viscometer and evaluated changes in morphology, gene expression, protein expression, and cytokine secretion over time. MDMs exposed to shear stress exhibited a rounder morphology compared to statically-cultured controls. RT-qPCR results showed significant upregulation of TNF-α, and analysis of cytokine release revealed increased secretion of IL-8, IL-18, fractalkine, and other chemokines. The upregulation of pro-inflammatory factors was evident with both increasing magnitudes of shear and time. Taken together, these results indicate that prolonged shear exposure induced a pro-inflammatory phenotype in human MDMs. These findings have implications for medical technology development, such as *in situ* vascular graft design wherein macrophages are exposed to shear and have been shown to affect graft resorption, and in delineating disease pathophysiology, for example to further illuminate the role of macrophages in atherosclerosis where shear is directly related to disease outcome.

## 1 Introduction

Macrophages are innate immune cells that are involved in wound healing, fighting infection, and stimulating other immune cells in response to injury. They readily adopt a wide spectrum of phenotypes regulated by the biochemical cues from their surrounding microenvironment, which enables these cells to play a critical role in wound healing regulation. Although macrophage polarization is a continuous spectrum *in vivo, in vitro* studies have simplified this spectrum into two distinct phenotypes: M1 and M2, with varying subsets [1, 2]. Upon exposure to pro-inflammatory signals such as interferon-γ (IFN-γ) and lipopolysaccharide (LPS), macrophages classically activate into the pro-inflammatory M1 phenotype and genetically express markers such as tumor necrosis factor-α (TNF-α) and chemokine receptor CCR7 [3–5]. M1 macrophages dominate the pro-inflammatory phase of wound healing, clearing cellular debris and secreting pro-inflammatory cytokines and chemokines, such as TNF-α, IL-1β, IL-6, IL-12, and IL-23 [6]. By contrast, exposure to anti-inflammatory signals such as interleukin (IL)-4 and IL-13 incurs alternative activation into the anti-inflammatory M2 phenotype characterized by the expression of transforming growth factor-β (TGF-β) and mannose receptor MRC1 [3, 4]. M2 macrophages are heavily involved in wound resolution and fibrosis and secrete factors such as IL-10 and TGF-β that dampen the inflammatory response [7]. In addition to differences in gene and protein expression, M1 and M2 macrophages display distinct morphological variations. M1 macrophages exhibit the traditional “fried egg” morphology, whereas M2 macrophages are elongated and spindle-like [8, 9]. Overall, it is accepted that biochemical signals induce complex responses from macrophages through changes in gene expression, protein secretion, and morphology.

While much of the current research has focused on the role of cytokines and chemokines in regulating macrophage plasticity, the role of the mechanical microenvironment has received considerably less attention. Macrophages respond to their surrounding mechanical microenvironment through a process called mechanotransduction. Through this process, they sense mechanical stimuli, such as stiffness and topography, which they convert into biochemical signals. Recent works have elucidated the pivotal role of mechanical cues in regulating macrophage differentiation and function, with the stiffness of the extracellular matrix being a key determinant [10–12]. Specifically, it has been demonstrated that increased matrix stiffness, characteristic of fibrotic tissues, drives macrophage polarization towards an anti-inflammatory M2 phenotype [12]. This finding is particularly significant given that M2 macrophages inherently exhibit a greater stiffness compared to their pro-inflammatory M1 counterparts [13]. This mechanical influence extends to morphological adaptations, with macrophages in fibrotic environments adopting an elongated shape [8].

Furthermore, manipulation of macrophage morphology itself has been shown to alter polarization states. McWhorter et al. demonstrated that artificially elongating macrophage morphology via engineered substrates enhances M2 marker expression while reducing pro-inflammatory cytokine production [8]. Conversely, Jain et al. observed that constraining the spread of inherently more spherical M1 macrophages dampens their pro-inflammatory activation [14]. Aside from stiffness and topology, macrophages can be influenced by dynamic mechanical loading, such as stretch and compression. Studies of human monocyte-derived macrophages (MDMs) have shown that cyclic stretch induces their expression of matrix metalloproteinase (MMP)-1 and MMP-3 and encourages their differentiation into osteoclasts for bone repair and maintenance [15, 16]. Overall, these investigations demonstrate that macrophage morphology and behavior are not only influenced by cellular signals, but also by the mechanical forces imposed upon them.

Although researchers have begun to investigate the effect of various mechanical stimuli on macrophages, there is a paucity of research over their response to shear stress. Recent works investigating macrophage behavior in shear stress conditions have primarily been focused on atherosclerotic plaque formation [6, 17, 18]. Specifically, *in vivo* mouse models of atherosclerosis have shown that steady high shear within straight regions of coronary arteries is atheroprotective, dominated by macrophages expressing M2 markers, and correlated with plaque stability, whereas disturbed low shear at bifurcations is atheroprone, rich in M1 macrophages, and vulnerable to plaque rupture [17, 19, 20]. However, there are other regions of the heart and cardiovascular system where the effects of high shear stresses are less clear. One example is the left ventricular outflow tract (LVOT), where high shear stresses are putatively associated with discrete subaortic stenosis (DSS), the formation of a membrane structure that encircles the LVOT and narrows the pathway to the aortic valve, increasing shear stress [21]. Analysis of septal myectomy in patients with DSS has revealed glycosaminoglycan and proteoglycan deposition, along with evidence of inflammation, suggesting that macrophages may contribute to the development of this fibrous membrane [22, 23].

The specific role of shear stress in macrophage response, however, remains an open question. Although other studies have evaluated hemodynamic loading on macrophages in the context of biomaterial-driven *in situ* vascular regeneration and shown that macrophage activation is augmented upon exposure to shear stress [24–26], there is limited knowledge about the biomolecules that macrophages release in response to these changes. Furthermore, *in vivo* studies have difficulty teasing out the specific effects of fluid shear stress on macrophages considering that other factors such as stiffness and cellular signaling also influence macrophage function. Therefore, in this study, we employed an *in vitro* model system to characterize macrophage behavior under various rates of physiological shear [27, 28]. Human MDMs were exposed to 15 dynes/cm^2^ and 35 dynes/cm^2^ shear stress over a period of 3, 24, and 48 hours using a cone-and-plate viscometer. The macrophages were then evaluated for morphology, gene expression, protein expression, and cytokine secretion, enabling the study of the direct effects of shear stress on macrophage behavior, which, in turn, can help better define the role of macrophages in cardiovascular diseases.

## 2 Materials and Methods

### 2.1 Primary human monocyte-derived macrophages

Human monocytes were isolated from donor buffy coats received from the Gulf Coast Regional Blood Center (Houston, TX). Use of de-identified blood product samples received through a third party were deemed by the Rice University IRB to not be human subjects’ research. The buffy coats were layered on top of 15 mL of Histopaque (1.077 g/mL; Sigma-Aldrich) and centrifuged at 400 x g for 30 minutes at room temperature without brake. Following centrifugation, the peripheral blood mononuclear cells (PBMCs) were collected from the interphase, pooled together from 3-4 donors, and washed twice with 1x PBS containing 5 mM EDTA. PBMCs were seeded onto non tissue-culture treated flasks in RPMI 1640 (Sigma-Aldrich) supplemented with 10% fetal bovine serum (FBS), 1% penicillin/streptomycin, and MCSF (20 ng/mL; Novus Biologicals). After 24 hours, the floating lymphocytes were aspirated, and the flask was washed 2x with PBS to remove remaining lymphocytes, leaving attached monocytes. Monocytes were cultured in a 5% CO2 incubator at 37°C for a subsequent 5-7 days in RPMI 1640 supplemented with 10% FBS, 1% penicillin/streptomycin, and MCSF (20 ng/mL). Media was replenished on days 3 and 5. On days 5-7, macrophages were harvested after incubation with Accutase (Sigma-Aldrich) for 15 minutes along with gentle cell scraping.

### 2.2 Shear stress application

We applied uniform laminar shear stress to primary human MDMs using a cone-and-plate viscometer system as previously described [29–31]. The cone was placed on top of a 60 mm petri dish seeded with a monolayer of MDMs that was then positioned on top of a controllable magnetic stir plate. The entire system was placed in a 5% CO_2_ incubator at 37°C. Each petri dish was washed with PBS and supplemented with 3 mL of fresh MDM media (RPMI 1640 supplemented with 10% FBS and 1% penicillin/streptomycin) prior to shear exposure. The cone was then rotated at 280 RPM or 660 RPM, which equated to a shear rate of 15 dynes/cm^2^ and 35 dynes/cm^2^, respectively, using the governing wall shear stress equation:

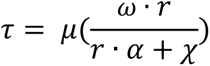

where τ is the desired shear stress, μ is the dynamic viscosity (0.95 centistokes) of the media, ω is the angular velocity of the cone, r is the radius (30 mm) of the cone, α is the angle (0.5°) of the cone, and *χ* is the gap height (300 µm) between the cone and the cells. MDMs were exposed to these shear rates for 3, 24, or 48 hours, and phenotypes were then evaluated through flow cytometry and RT-qPCR. The 48-hour 35 dynes/cm^2^ group was supplemented with an extra 1 mL of RPMI 1640 without serum after 24 hours to prevent the samples from drying out during the duration of the experiment. The addition of serum-free media resulted in a consistent final volume between all groups.

### 2.3 Flow cytometry

After isolation and differentiation, MDMs were evaluated for purity through flow cytometry for expression of the pan-macrophage marker, CD68. MDMs were harvested after 15 minutes of Accutase exposure through gentle scraping. For intracellular staining, MDMs were fixed with 4% paraformaldehyde for 20 minutes at 4°C, washed 2x with 1% FBS in PBS, and permeabilized in PBS with 0.5% Tween20 and 1% FBS for 15 minutes at room temperature. Cells were then washed 2x with 1% FBS in PBS and incubated in 1% FBS in PBS with APC-conjugated mouse anti-human CD68 (BD Biosciences) for 1 hour at room temperature. Following staining, samples were washed 2x with 1% FBS in PBS and transferred to 5 mL flow cytometry tubes.

Following shear stress application, the MDM protein expression of characteristic phenotypic markers was evaluated using flow cytometry. Cells were stained with the following antibodies: FITC-conjugated mouse anti-human CD80 (BD Biosciences), PE-conjugated mouse anti-human CD163 (BD Biosciences), and APC-conjugated mouse anti-human CD206 (BD Biosciences). Briefly, MDMs were harvested after 15 minutes of Accutase exposure through gentle scraping. Cells were incubated in 1% FBS in PBS with all antibodies for 30 minutes at room temperature. Non-specific binding was blocked by incubating the samples with 1% FBS in PBS. Following staining, samples were washed 2x with 1% FBS in PBS and transferred to 5 mL flow cytometry tubes. Flow cytometry data for all markers was acquired using the Sony MA900 multi-application cell sorter (Sony Biotechnology Inc) and was analyzed using FlowJo.

### 2.4 Viability staining and morphology quantification

Cell viability was assessed following shear stress application by incubating each sample in a solution containing 2 µM calcein AM and 4 µM ethidium homodimer-1 in PBS (Live/Dead Viability/Cytotoxicity Kit; Thermo Fisher) for 30 minutes at room temperature. Samples were imaged immediately via fluorescent microscopy. Cell viability was quantified through FIJI. Morphology was analyzed using FIJI’s circularity feature where circularity is defined as 4π*(area/perimeter^2^). A perfect circle has a circularity of 1. The circularity of each individual cell was analyzed from each group’s live/dead images resulting in a large number of samples.

### 2.5 RNA isolation, cDNA synthesis, and quantitative reverse transcription PCR

Following shear stress application, MDM phenotypes were evaluated through quantitative reverse transcription PCR (RT-qPCR). RNA was extracted using the Direct-zol™ RNA Microprep kit (Zymo Research) according to the manufacturer’s instructions. Briefly, MDMs were lysed with TRIzol (Thermo Fisher), mixed with an equal volume of 95% ethanol, and transferred to a Zymo-Spin™ IC Column in a collection tube. Samples were then incubated with DNase I for 15 minutes at room temperature, washed with RNA wash buffer, and eluted with 15 µL of DNase/RNase-free water. RNA purity (A_260/280_ > 1.8) was then assessed through the NanoDrop 2000 spectrophotometer (Thermo Fisher), and RNA was stored at −80°C until further steps. Complementary DNA (cDNA) was synthesized from purified RNA using the High-Capacity cDNA Reverse Transcription Kit (Applied Biosystems) according to manufacturer’s instructions and then stored at −20°C until gene expression analysis. RT-qPCR was performed using the iTaq™ Universal Probes Supermix (Bio-Rad Laboratories) according to manufacturer’s instructions with primers specific to human CCR7 (probe ID: Hs01013469_m1), MRC1 (Hs02832368_g1), TGFB1 (Hs00998133_m1), TNF-α (Hs00174128_m1), and GAPDH (Hs02786624_g1) (Applied Biosystems). Fold changes were first normalized to GAPDH and then to the control group (no shear stress application).

### 2.6 Conditioned media analysis

After shear exposure, conditioned media containing cytokines secreted by MDMs were collected and centrifuged at 1000 x g for 5 minutes to remove cellular debris. The supernatant was then collected and stored at −80°C until cytokine analysis. The supernatants were from the same samples that were analyzed through RT-qPCR so that a direct comparison between gene expression and protein secretion could be made. Cytokines were analyzed using the MILLIPLEX® Human 48-Plex Cytokine Panel A kit (MilliporeSigma) on the Luminex system as per manufacturer’s instructions. This analysis was done by the proteomics core laboratory at Baylor College of Medicine, Houston, TX. Briefly, samples were thawed on ice and centrifuged at 10000 x g for 5 mins. Samples and beads were incubated at 4°C on a shaker for 16 hours. Well contents were then removed, and the plate was washed 3x with 200 µL Wash Buffer. 25 µL of detection antibodies were added to each well and incubated at room temperature for 1 hour. 25 µL of Streptavidin-Phycoerythrin was then added to each well containing the detection antibodies and incubated for 30 minutes at room temperature. Well contents were removed, and the plate was washed 3x with 200 µL Wash Buffer. 150 µL of Sheath Fluid PLUS was added to all wells, beads were resuspended on a plate shaker for 5 minutes, and the plate was read on the Luminex system. RPMI-1640 media was used as the blank. Two quality control samples were prepared, and a Logistic-5PL regression type was used to construct the standard curve of all the analytes.

### 2.7 Statistical analysis

Statistical analysis was performed using GraphPad Prism software version 6 (GraphPad Software Inc.) and R. A two-way or one-way analysis of variance (ANOVA) was used to determine statistical significance (p < 0.05) for each set of conditions in this study. P-values were adjusted for multiple comparisons using post hoc Tukey’s testing. All experiments were repeated in triplicates at a minimum.

## 3 Results

### 3.1 Shear stress exposure modulated MDM morphology by reducing elongation

A cone-and-plate viscometer was used to apply varying shear rates (0 dynes/cm^2^ (static), 15 dynes/cm^2^, and 35 dynes/cm^2^) to MDMs for a period of 3, 24, and 48 hours (**Fig. 1**). Post shear exposure, samples were treated with live/dead stain and imaged to evaluate cell viability (**Fig. 2A**). As shown in the fluorescent images, cell viability did not change over time or shear stress (93% average viability, **Fig. 2B**). Additional bright-field images were taken to examine cell morphology (**Fig. 3**). Morphological changes were determined using the circularity measurement in FIJI. Interestingly, all groups were generally more elongated (less circular) as time increased (**Fig. 3, Supplemental Fig. 1**). Exposure to shear at each timepoint increased circularity, aside from the 48-hour timepoint where the 35 dynes/cm^2^ condition was not significantly different than static. These results indicate that MDMs were expressing a rounder morphology in response to shear.

**Fig. 1.**
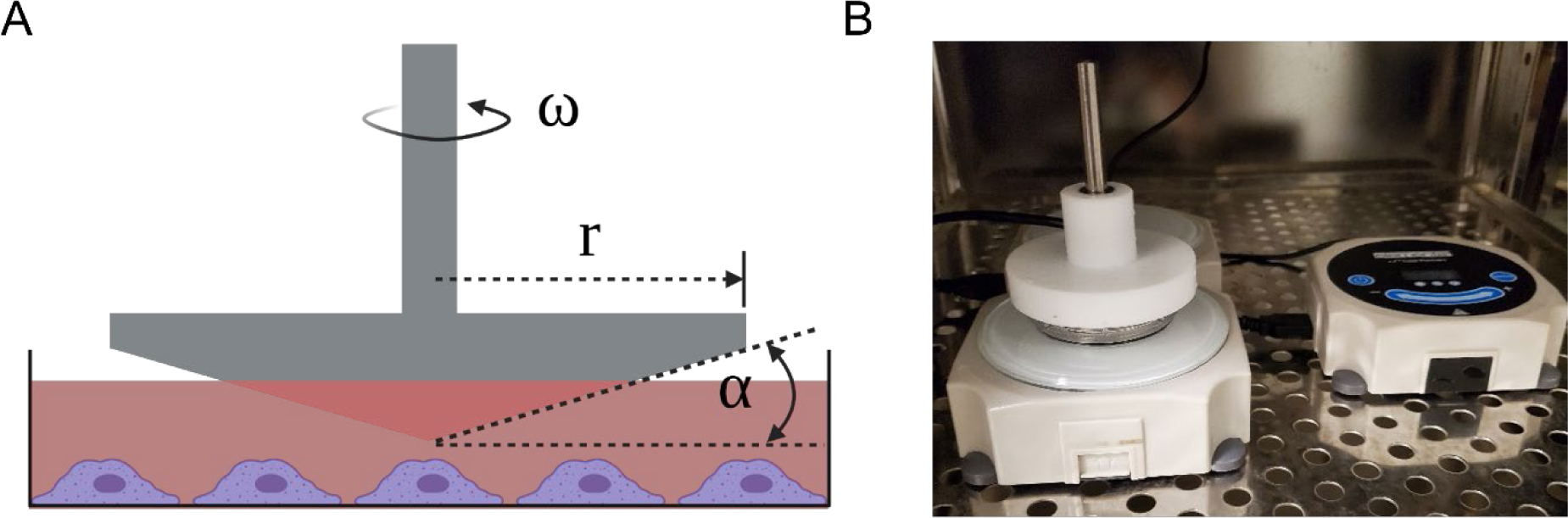
Cone-and-plate system used to apply shear stress to MDMs. (A) Schematic of the cone-and-plate system where MDMs are attached to the bottom of the petri dish and a cone is loaded on top. (B) Setup of the cone-and-plate system in the incubator. The right side shows the controller that sets the RPM of the system

**Fig. 2.**
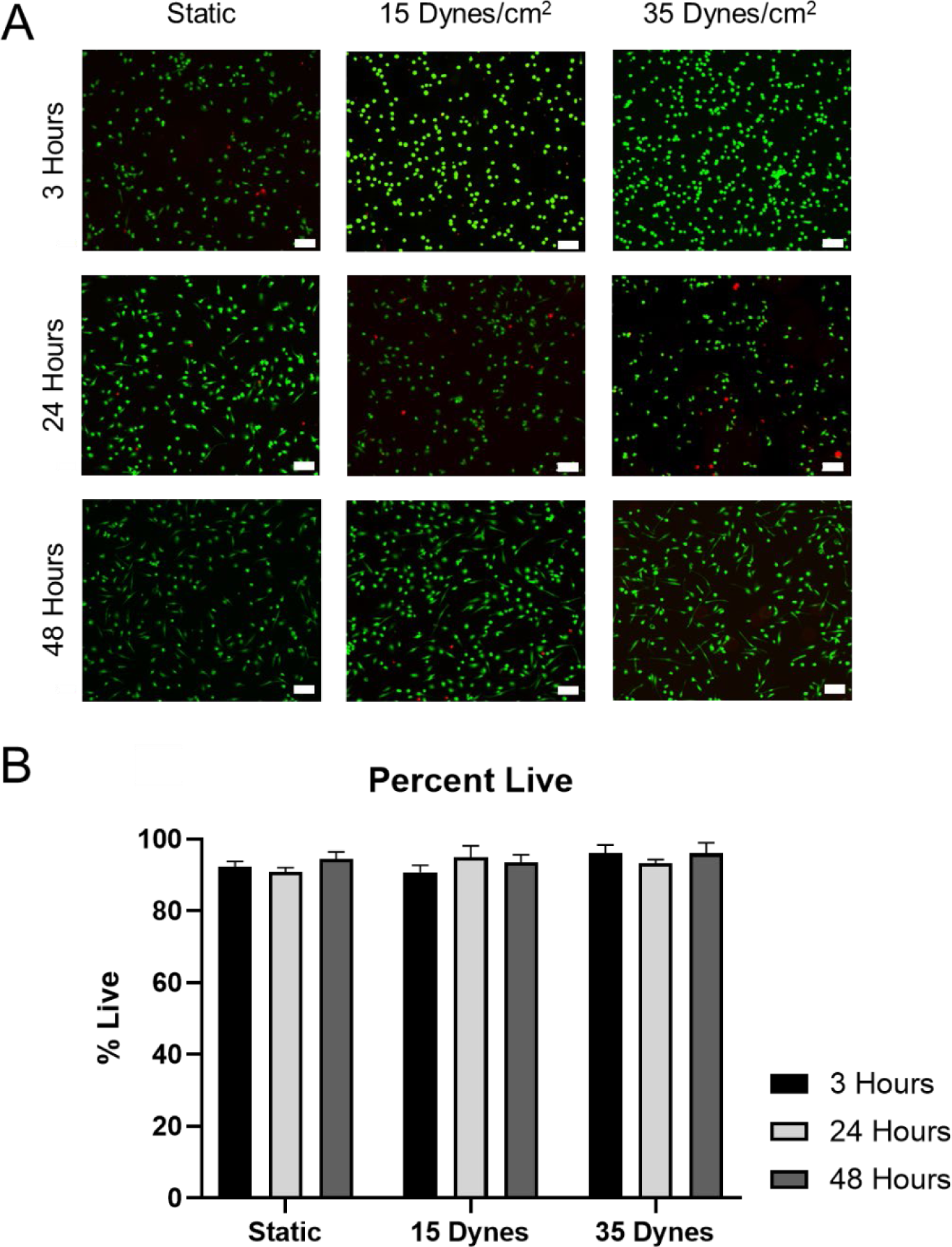
MDM viability following shear exposure. (A) Live/dead staining and (B) quantification of MDM viability was performed following exposure to static, 15 dynes/cm^2^, and 35 dynes/cm^2^ conditions for a period of 3, 24, and 48 hours. Each image was taken immediately after shear exposure at 3, 24, and 48 hours and is representative of all data collected. Scale bar = 100 µm

**Fig. 3.**
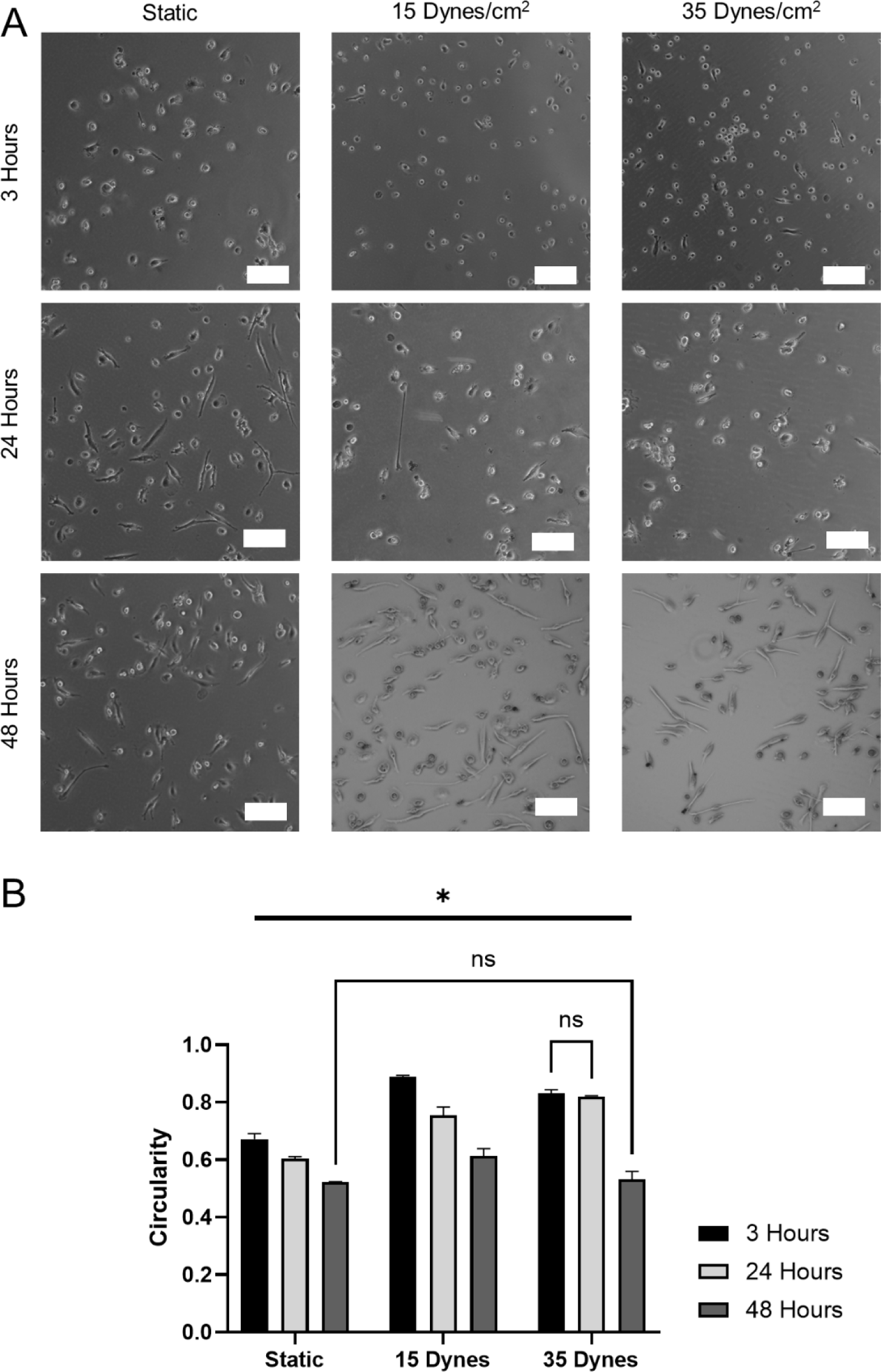
MDM morphology following shear exposure. (A) Brightfield imaging and (B) circularity analysis of MDM morphology was performed following exposure to static, 15 dynes/cm^2^, and 35 dynes/cm^2^ conditions for a period of 3, 24, and 48 hours. Scale bar = 100 µm. Data is represented as mean ± SD, n = 3. *Represents all pairwise comparisons are statistically significant at p < 0.05 except those indicated by ns

### 3.2 High shear stress attenuated CD80, CD163, and CD206 protein expression

MDM protein expression was also evaluated through flow cytometry after shear exposure (**Fig. 4**). We investigated CD80 as a marker for a pro-inflammatory phenotype, and CD206 and CD163 as markers for an anti-inflammatory phenotype. CD80 surface marker expression was downregulated in response to increasing time in all groups. This downregulation was evident across all timepoints in response to increasing shear except for 48 hours where expression remained unchanged. Interestingly, CD163 expression generally remained unchanged in the static condition over time after an initial downregulation but exhibited attenuated expression levels in response to shear. Similarly, CD206 expression remained unaffected by time within the static and 15 dynes/cm^2^ groups but was significantly downregulated at 48 hours in the 35 dynes/cm^2^ group. Overall, prolonged exposure to increased levels of shear stress reduced CD80, CD163, and CD206 protein expression, demonstrating a mechanosensitive response.

**Fig. 4.**
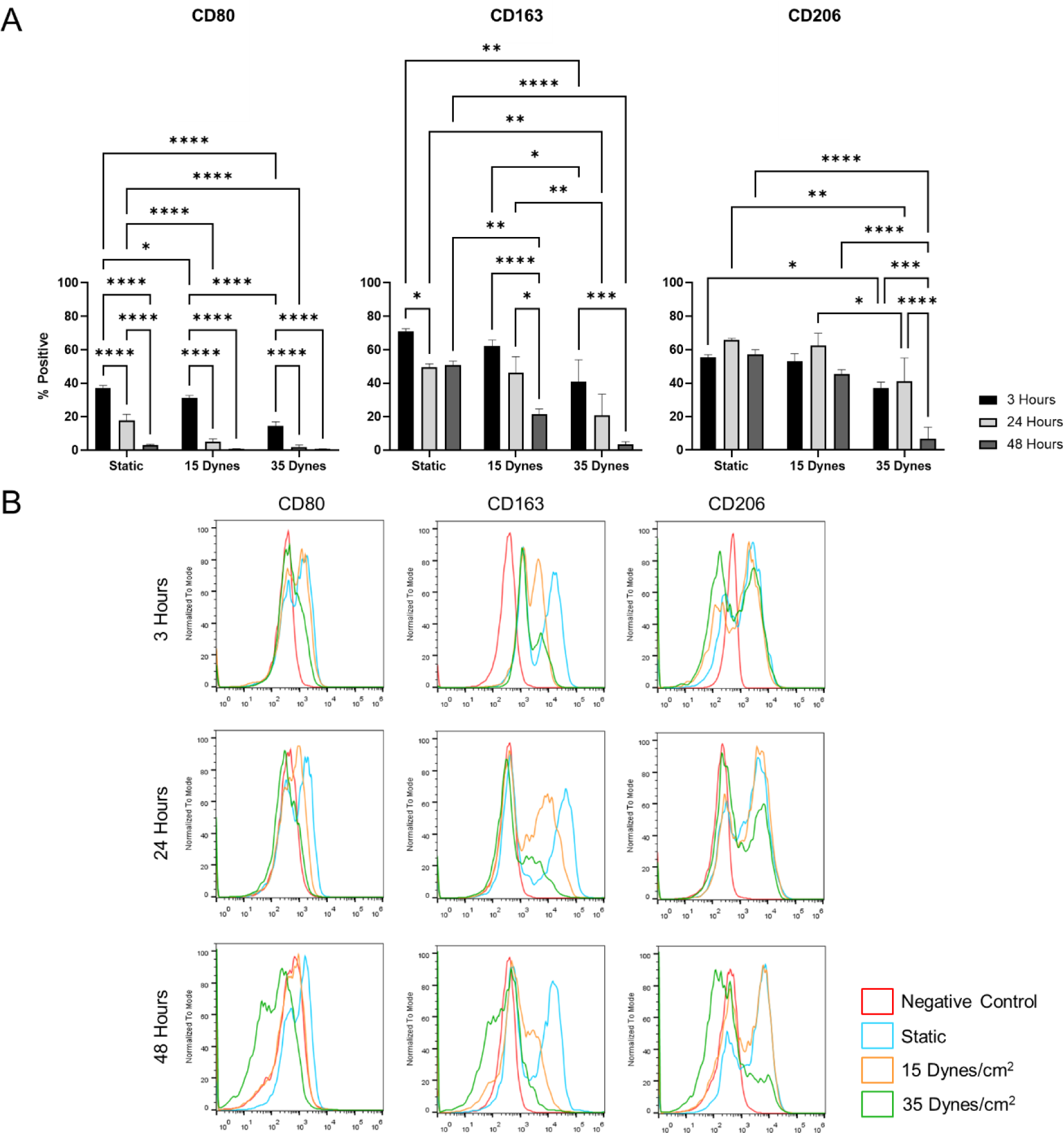
MDM protein expression following shear exposure. (A) Flow cytometry was performed to quantify percent positive protein expression of CD80, CD163, and CD206 following shear exposure. (B) Representative histograms showing the fluorescent signal intensity of each cell. Data is represented as mean ± SD, n ≥ 3

### 3.3 Shear stress upregulated gene expression of TNF-α in a time-dependent manner

After shear exposure, RNA was isolated from samples, and RT-qPCR was performed to evaluate MDM gene expression. We investigated TNF-α and CCR7 as markers for a pro-inflammatory phenotype; and TGF-β1 and MRC1, which codes for CD206, as markers for an anti-inflammatory phenotype. TNF-α was initially downregulated in response to shear exposure but became most elevated at the 24-hour timepoint where increasing magnitudes of shear promoted significant upregulation compared to static controls (7.36 vs. 16.87-fold change for the two shear magnitudes, respectively, **Fig. 5**). At 48 hours, TNF-α was upregulated when exposed to 15 dynes/cm^2^ of shear but remained unchanged in response to 35 dynes/cm^2^ of shear, suggesting a time-dependent response. TGF-β1, MRC-1, and CCR7 exhibited no significant response at 3 hours, however at both 24 and 48 hours, these genes were downregulated in response to monotonically increasing shear. Overall, the prominent upregulation of TNF-α in response to shear within the 24-hour timepoint indicated that MDMs may be expressing a time-dependent inflammatory phenotype upon exposure to shear.

**Fig. 5.**
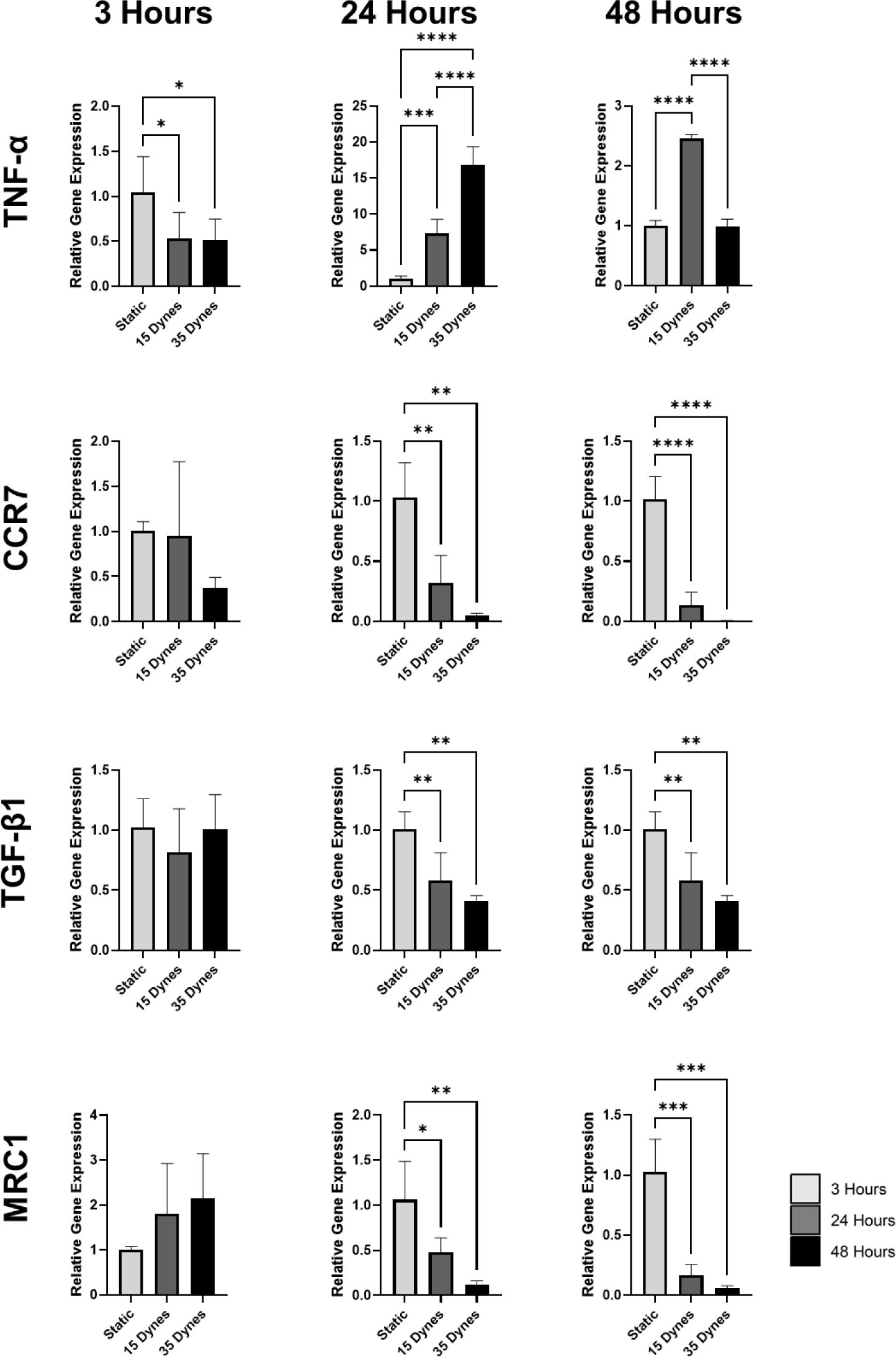
MDM gene expression following shear exposure. RT-qPCR was performed to evaluate gene expression of TNF-α, CCR7, TGF-β1, and MRC1 from MDMs cultured under static, 15 dynes/cm^2^, and 35 dynes/cm^2^ conditions for a period of 3, 24, and 48 hours. Data is represented as mean ± SD, n ≥ 3. *p < 0.05, **p < 0.01, ***p < 0.001, ****p < 0.0001

### 3.4 Shear stress promoted time-dependent pro-inflammatory cytokine secretion

#### 3.4.1 Cytokine secretion in response to time within each shear condition

Since gene expression of TNF-α was most prominently upregulated, we analyzed conditioned media for cytokine levels using a 48-plex cytokine panel. The results shown in **Figs. 6-8** demonstrate that both shear stress and time affected cellular signaling. Interestingly, within each shear condition, increasing time overwhelmingly correlated with monotonically increased production of chemokines and pro-inflammatory cytokines (**Figs. 6-7**). IFN-γ, granulocyte-colony stimulating factor (G-CSF), and numerous chemokines (IL-8, IL-15, monocyte chemoattractant protein-1 (MCP-1), macrophage-derived chemokine (MDC), and chemokine ligand 9 (CXCL9)), were all increased in response to prolonged time across the different shear conditions. In contrast, the inflammatory chemokines, macrophage inflammatory protein (MIP)-1α and MIP-1β were both downregulated over time, but only in the static condition.

**Fig. 6.**
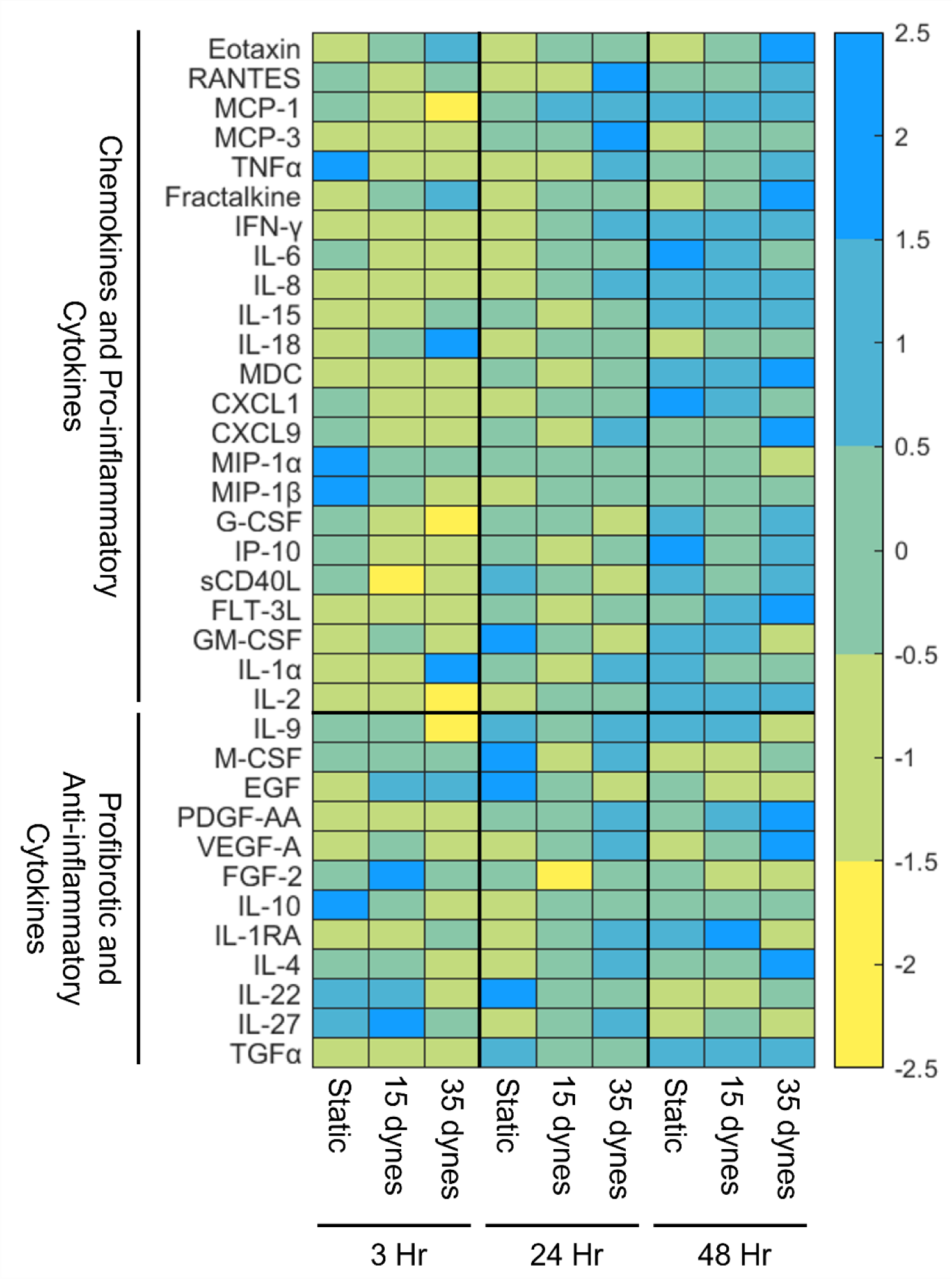
Heatmap of cytokines released by MDMs cultured under static, 15 dynes/cm^2^, and 35 dynes/cm^2^ conditions for a period of 3, 24, and 48 hours. Visualizes inflammatory cytokine secretion as time and shear increases. Cytokine secretion level is represented using z-scores

**Fig. 7.**
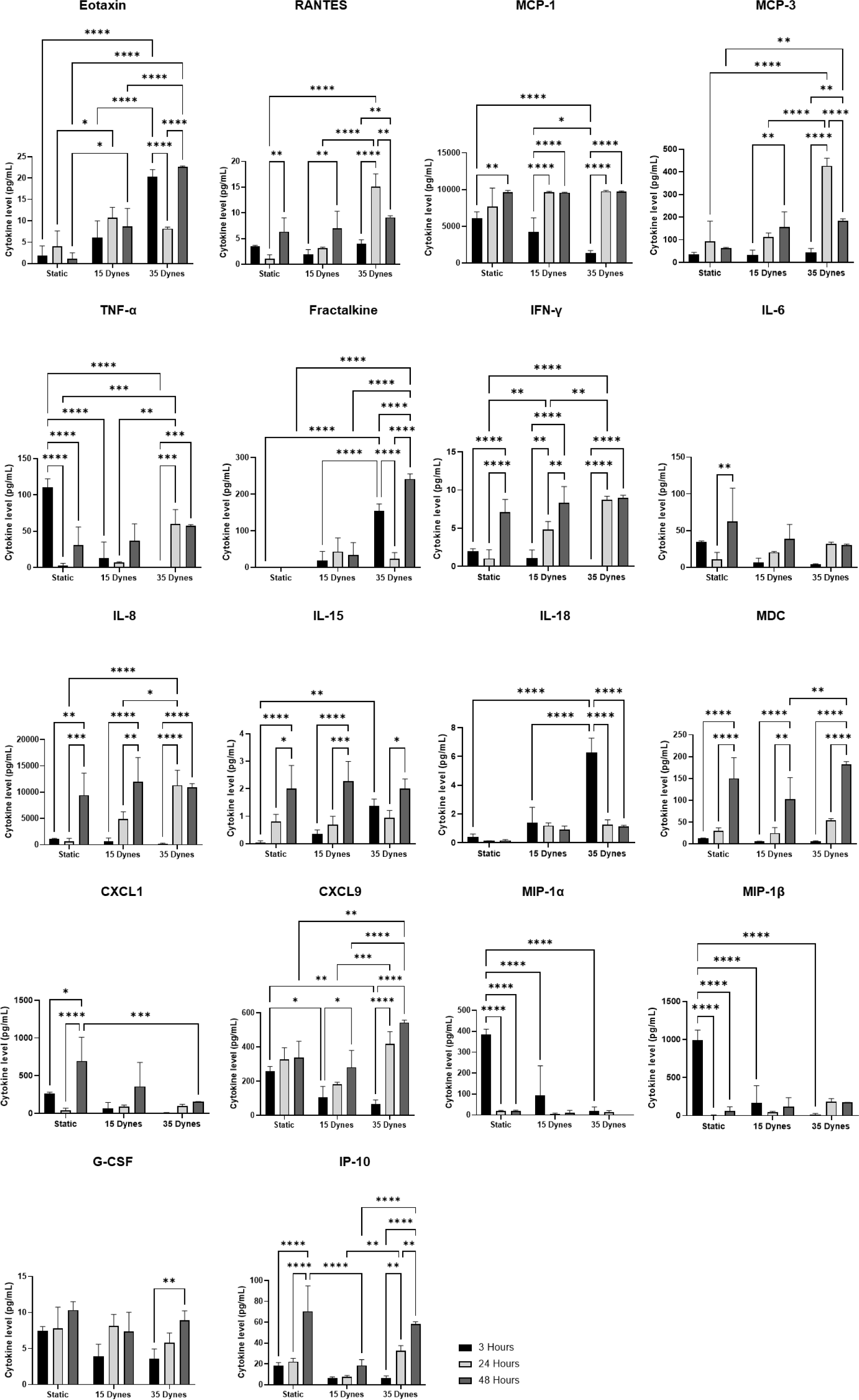
Chemokines and pro-inflammatory cytokine secretion from MDMs cultured under static, 15 dynes/cm^2^, and 35 dynes/cm^2^ conditions for a period of 3, 24, and 48 hours. Data is expressed as mean ± SD, n = 4. *p < 0.05, **p < 0.01, ***p < 0.001, ****p < 0.0001

**Fig. 8.**
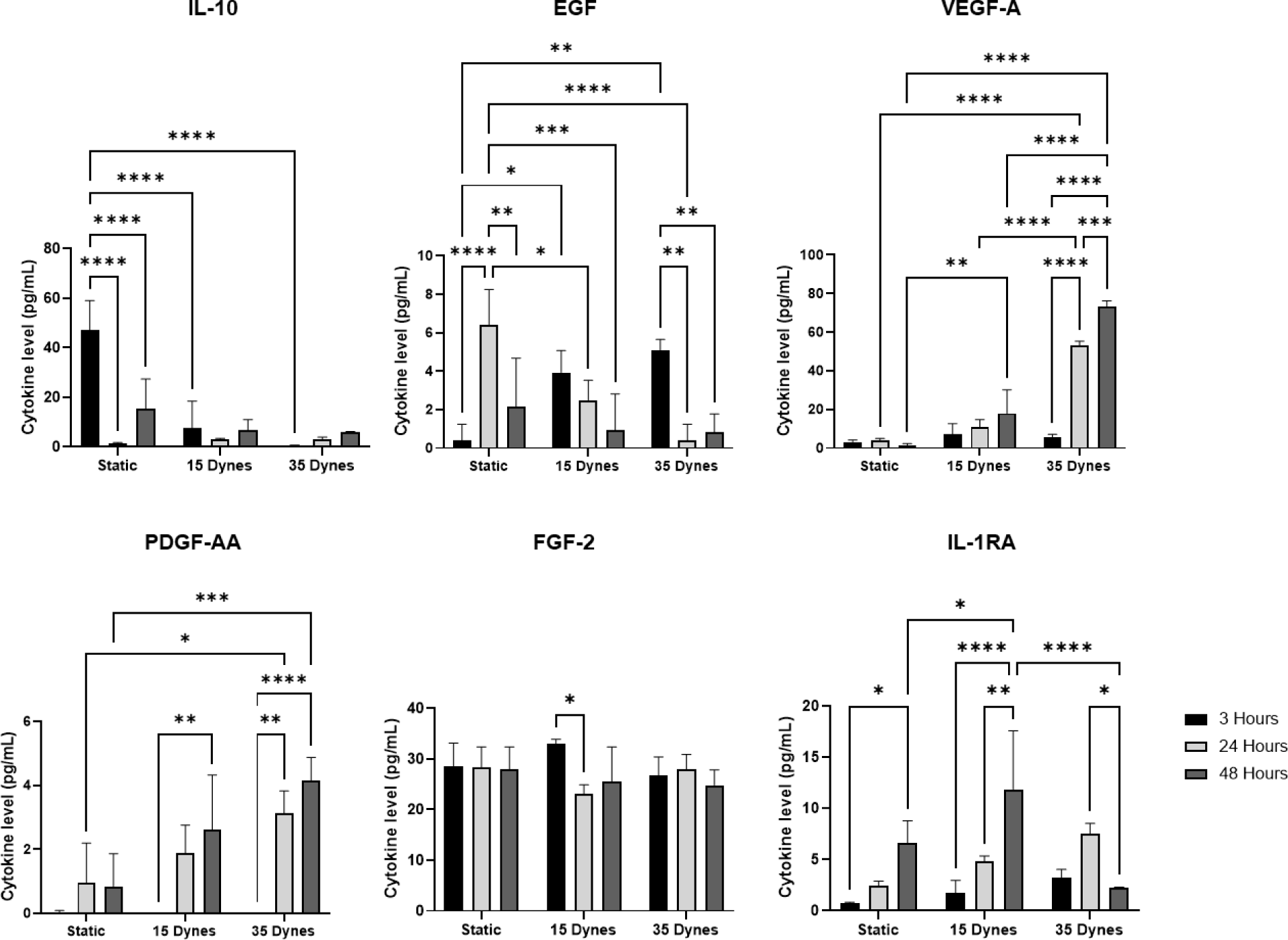
Anti-inflammatory cytokine secretion from MDMs cultured under static, 15 dynes/cm^2^, and 35 dynes/cm^2^ conditions for a period of 3, 24, and 48 hours. Data is expressed as mean ± SD, n = 4. *p < 0.05, **p < 0.01, ***p < 0.001, ****p < 0.0001

Secretion of the anti-inflammatory cytokines, however, was varied (**Figs. 6&8**). Epidermal growth factor (EGF) production in the static condition was upregulated at 24 hours (vs. 3 hours) but was downregulated at 48 hours. Furthermore, it remained significantly downregulated in the 15 and 35 dynes/cm^2^ conditions as time increased. Fibrotic growth factor-2 (FGF-2) remained generally unchanged over time but had decreased levels at 24 hours vs. 3 hours in the 15 dynes/cm^2^ condition. IL-1RA was upregulated across all timepoints within the static and 15 dynes/cm^2^ conditions but was downregulated at 48 hours in the 35 dynes/cm^2^ condition. Vascular endothelial growth factor-A (VEGF-A) and platelet derived growth factor-AA (PDGF-AA) were upregulated in the 15 and 35 dynes/cm^2^ conditions over time but remained unchanged in the static condition.

#### 3.4.2 Cytokine secretion in response to increasing shear magnitude within each timepoint

Increasing shear, however, stimulated paradoxical effects within different timepoints. Among the pro-inflammatory cytokines (**Figs. 6-7**), CXCL9 and TNF-α were downregulated in response to increasing shear at 3 hours. However, at 24 and 48 hours, these cytokines were upregulated following increased shear. Secretion of IFN-γ and IL-8 was unaffected at 3 hours but was significantly upregulated in response to shear at 24 hours. MCP-3 secretion was unaffected by shear at 3 hours but was upregulated at 24 and 48 hours in response to 35 dynes/cm^2^ of shear. MCP-1 secretion initially decreased at 3 hours in response to increasing shear but remained unaffected at 24 and 48 hours. RANTES was not affected by increased shear at 3 and 48 hours but was significantly upregulated in response to 35 dynes/cm^2^ of shear at 24 hours. MIP-1α and MIP-1β were significantly downregulated at 3 hours with increasing shear but were unaffected at 24 and 48 hours. Eotaxin and fractalkine/CX_3_CL1, potent chemoattractants for eosinophils and monocytes/T cells, respectively, showed more secretion with increased shear at every timepoint. However, this result was most significant at the 3-hour and 48-hour timepoints. Interestingly, fractalkine was not found in the static conditioned media at any timepoint.

Looking at the effects of increased shear on anti-inflammatory cytokine release within each timepoint (**Figs. 6&8**), EGF was upregulated in response to increased shear at 3 hours. However, the 24-hour timepoint showed the opposite pattern with less EGF secretion in response to increased shear. IL-10, a major cytokine secreted by M2 macrophages, was profoundly downregulated following increased shear at 3 hours, and maintained low abundance for all shear conditions at the 24- and 48-hour timepoints. PDGF-AA and VEGF-A secretions were unchanged by shear at the 3-hour timepoint but were monotonically upregulated by increasing shear at both the 24- and 48-hour timepoints. IL-1RA secretion was unchanged by shear at the 3- and 24-hour timepoints but was upregulated at 48 hours in the 15 dynes/cm^2^ condition. Overall, the cytokine panel revealed a predominantly pro-inflammatory response towards prolonged shear exposure.

## 4 Discussion

Macrophages are highly plastic cells that exhibit diverse phenotypes in response to changes in their microenvironment, which enable them to play crucial roles in the wound healing process; correspondingly, they are heavily involved in major disease outcomes [23, 32–34]. Although recent investigations have focused on the response of macrophages to the mechanical aspects of their environment, there is limited research surrounding macrophage behavior under shear stress. In the present study, we subjected human MDMs to physiological levels of shear using a cone-and-plate viscometer over the course of 3, 24, and 48 hours. Our main findings were that MDMs expressed time-dependent changes in pro-inflammatory cytokine secretion, gene expression, and morphology in response to shear stress. Specifically, initial exposure to shear stress downregulated pro-inflammatory cytokine secretion at 3 hours, but an opposite effect was observed after prolonged shear at 24 and 48 hours. This phenotypic switch was also consistent when looking at gene expression, where TNF-α was upregulated while TGF-β1 and MRC1 were downregulated at 24 hours.

Evaluation of gene expression and morphology supports that MDMs express a pro-inflammatory phenotype after shear exposure. MDMs in the static condition became more elongated over time, suggesting that they were becoming more anti-inflammatory in normal culture. However, the application of shear curbed their elongation and promoted formation of a round morphology, suggesting a more pro-inflammatory state (**Supplemental Fig. 1**). Although there was a shift in MDM morphology indicative of a phenotypic switch, protein expression for both phenotypes were attenuated with shear exposure. The heterogeneity of protein expression observed could be attributed to the pooling of donors although all cells were positive for pan-macrophage marker CD68 (**Supplemental Fig. 2**). The downregulation of CD80 expression over time in the static condition is consistent with the morphology results, indicating that MDMs are becoming less inflammatory over time without shear exposure. Studies have also shown that macrophages change their phenotype over time in normal culture, exhibiting a transient pro-inflammatory profile [35, 36]. However, we expected the application of shear stress to promote CD80 expression. This attenuation could be due to macrophages expressing other pro-inflammatory markers that were not investigated such as CD86. The predominant findings for gene expression were that anti-inflammatory markers TGF-β1 and MRC1 were significantly downregulated in response to shear. The downregulation of CCR7 was unexpected, but consistent with previous similar works [26]. However, TNF-α, a prominent pro-inflammatory marker, was significantly upregulated at 24 hours by a large magnitude. Overall, the reduction in MDM elongation coupled with the significant upregulation of TNF-α gene expression indicated that the MDMs may be primed towards a pro-inflammatory phenotype following shear exposure, prompting further evaluation of cytokines.

The application of shear stress, over time, largely induced pro-inflammatory cytokine secretion by human MDMs. This finding was demonstrated by noting that protein levels of IFN-γ, a potent activator of pro-inflammatory macrophages, G-CSF, a neutrophil regulator, and numerous chemokines (eotaxin, fractalkine, IL-8, IL-15, MCP-1, MCP-3, MDC, RANTES, and CXCL9), significantly increased in response to prolonged shear exposure. These chemokines potently attract a wide variety of leukocytes including neutrophils, T-cells, monocytes, and macrophages [37–44]. Notably, fractalkine was not expressed in the static condition at any timepoint, indicating that shear stress exposure definitively induces macrophage-derived fractalkine secretion. Interestingly, protein levels for chemokines MIP-1α and MIP-1β were significantly downregulated with shear exposure. The conflicting results between chemokine expressions over time could be the result of different mechanisms of regulation. The mechanisms promoting MIP-1α expression in macrophages are not well understood, although the TLR4-IRF3 pathway is reported to be involved [45]. Furthermore, MIP-1α release by macrophages is reported to be downregulated by IFN-γ *in vitro*, which is consistent with our results [46]. IL-10, which is distinctly produced by anti-inflammatory macrophages, was heavily suppressed with increasing shear. Interestingly, VEGF-A and PDGF-AA had higher levels of expression over longer time points in both shear conditions. Both growth factors are heavily regulated by anti-inflammatory macrophages and involved in cell differentiation, proliferation, and angiogenesis. However, VEGF-A has also been reported to stimulate monocyte/macrophage chemotaxis [47]. Furthermore, studies have also shown that pro-inflammatory macrophages secrete the highest levels of VEGF-A compared to other macrophage phenotypes, and that IL-10 suppresses M1 but not M2-derived VEGF, which is consistent with our data [5, 48]. These results suggest that VEGF-A plays a pro-inflammatory role in response to shear stress. Overall, the overwhelming upregulation of chemokines suggests that macrophages are becoming more pro-inflammatory and releasing signals to attract other leukocytes under prolonged exposure to shear stress.

Although the end result of MDM shear exposure was inflammation, an interesting phenomenon occurred at the 3- and 24-hour timepoint. Several pro-inflammatory cytokines, including TNF-α, IFN-γ, IL-8, MCP-1, MCP-3, and RANTES, were initially downregulated or unchanged in response to shear with subsequent upregulation at 24 hours. However, EGF, a predominantly anti-inflammatory cytokine, exhibited an opposite effect with initial upregulation at 3 hours and subsequent downregulation at later timepoints, suggesting that EGF may play a role in regulating downstream pro-inflammatory macrophage response. The stark dichotomy between pro-inflammatory cytokine expression at the 3-hour timepoint compared to EGF is worth investigating in the context of atherosclerosis, since these cytokines have all been implicated in the initiation and progression of atherosclerotic plaques. The interaction between RANTES and fractalkine is established as an early pathway for rolling monocytes to firmly adhere to activated endothelium, and MCP-1 enables transendothelial migration [49]. IL-8 assists with the firm adherence of monocytes to endothelium but is also implicated in potentiating plaque angiogenesis with its elevated presence in late-stage lesions [49]. Furthermore, CXCL9 is known to promote differentiation and multiplication of leukocytes, cause tissue extravasation, and is upregulated in atherosclerosis [37, 50]. Studies have also shown that the inhibition of TNF-α and IFN-γ have reduced atherosclerotic lesion size and regressed plaque progression [51, 52]. Additionally, in atherosclerosis, endothelial denudation can occur, leading to plaque erosion and exposure of the underlying macrophages to shear stress [53–56]. The mechanisms promoting plaque erosion are not well understood, however it has been shown that eroded plaques expressed elevated levels of EGF compared to ruptured plaques [57, 58]. Interestingly, inhibition of EGFR has been demonstrated to attenuate atherosclerosis by decreasing pro-inflammatory cytokines, such as TNF-α and IL-6 [59, 60].

Since EGF is more elevated in eroded plaques compared to ruptured plaques, our results speculate that continued elevated cytokine production by MDMs could lead to eventual atherosclerotic plaque rupture. Given that our results show an initial increase in EGF secretion followed by subsequent downregulation, an interesting question postulates whether shear stress enhances the activity of EGFR, promoting the downstream pro-inflammatory response. One study observed an increase in EGFR expression and activation in response to fluid flow in a 3D ovarian cancer model [61]. Interestingly, mutant EGFR, along with Ptpn11, which encodes for protein-tyrosine-phosphatase Shp2, has also been implicated in the development of defective semilunar cardiac valves causing enlargement from an overabundance of mesenchymal cells [62–64]. Specifically, activation of the MAPK pathway through EGFR has been proven to drive epithelial–mesenchymal transition (EMT) in semilunar valve formation [64, 65]. In the context of discrete subaortic stenosis, where altered shear and circulating immune cells are hypothesized to play a role in subvalvular fibrosis [21], it would be interesting to investigate the interactions between MDMs, shear stress, and EGFR.

The results from our studies should also be considered when designing *in situ* vascular grafts. Acellular vascular implants have been a promising approach for replacing small-diameter arteries. These biodegradable scaffolds promote new tissue formation, cellular infiltration, and ECM deposition [66]. However, the rate of scaffold degradation heavily influences patient outcomes. If the degradation is too quick, insufficient repair and loss of structural integrity occurs [67]. In contrast, slow degradation could cause chronic inflammation and subsequent fibrosis [67]. Macrophages have been demonstrated to affect the rate of scaffold degradation given their secretion of cytokines promoting both inflammation and wound resolution [26, 68]. Consequently, the design of scaffolds that modulate the balance between pro-inflammatory/anti-inflammatory phenotypes has been a promising approach. Given that these scaffolds are exposed to shear stress after implantation, it is important to consider its effect on MDMs. Elevated cytokine production by MDMs would increase the rate of scaffold degradation, which should be accounted for when designing these grafts.

To our knowledge, this is the first study to characterize MDM phenotype in response to shear stress *in vitro*. While our results demonstrating the effect of shear stress on MDMs are very compelling, they must be considered in context. Although donors were kept consistent for RT-qPCR and Luminex analysis, a different set of donors was required for the flow cytometry analysis due to cell shortage. This could possibly account for the conflicting results between these data. Furthermore, it is difficult to test longer timepoints with this model without media replenishment given that media evaporation will occur at longer timepoints. Although cells were seeded at a consistent density, some cell detachment after shear exposure is expected and could contribute to varying expression levels. Nonetheless, the results of gene expression, protein secretion, and morphology all demonstrate that exposure to shear stress drives a pro-inflammatory response from human MDMs, which is consistent with previous similar works [17, 56].

Future investigations should extend beyond our current study’s timeframe to include timepoints up to 72 hours to understand the temporal dynamics of macrophage responses to shear stress more comprehensively. A focused examination on the mechanisms driving the pro-inflammatory phenotype should involve targeted inhibition of the NF-κB and MAPK pathways using specific inhibitors like PS-1145 and SB203580, respectively, to delineate their contributions during shear stress and re-evaluate macrophage phenotype [69, 70]. Furthermore, while our study focused on the protein expression changes, we recognize the interconnected nature of cellular responses, which prompts us to consider the potential impact on other cellular components such as metabolites. Recent studies, such as those discussed by Evers et al., highlight the established link between mechanical cues and cellular metabolism, suggesting that shear stress may not only influence protein expression but also induce significant metabolic changes [71]. Mechanotransducers such as YAP/TAZ have been shown to regulate metabolic changes in many cell types. In mesenchymal stem cells, shear stress enhances YAP expression, driving their differentiation into chondrocytes [72]. In light of our findings, a promising direction for future research would be to investigate how shear stress affects the YAP/TAZ signaling pathway in macrophages and its subsequent impact on metabolic responses. It has previously been demonstrated that YAP/TAZ directly promotes a pro-inflammatory metabolic response by modulating IL-6 and Arginase-I expression via the histone deacetylase 3 (HDAC3)-nuclear receptor corepressor 1 (NCoR1) repressor complex [73]. Such studies will not only deepen our understanding of macrophage biology under mechanical stress but also may reveal new therapeutic targets for modulating inflammation.

In conclusion, in the present study, we explored the response of human MDMs to shear stress, revealing a time-dependent shift toward a pro-inflammatory phenotype characterized by the upregulation of pro-inflammatory cytokine secretion, gene expression, and cellular morphology. Our findings highlight the complex interplay between mechanical stimuli and macrophage activation, with significant implications for understanding atherosclerotic plaque dynamics and the design of vascular grafts. This research not only contributes to our knowledge of macrophage biology under shear stress but also highlights the potential for targeted interventions to modulate inflammatory responses in vascular pathologies and biomaterial design.

## 6 Author contributions

Conceptualization: EJ, JC, SK, and JG-A. Investigation: EJ, GK, and HP. Formal analysis: EJ. Writing – original draft: EJ. Writing – review & editing: EJ, GK, KS, JP, and JG-A. Methodology: EJ, GK, KS, JP, and JG-A. Supervision: RB, JP, SK, and JG-A.

### Funding acquisition

SK and JG-A. All authors contributed to the article and approved the final submitted version.

## 7 Data availability statement

The datasets generated for this study are available on request from the corresponding author.

## Acknowledgements

The authors would like to thank Dr. Shixia Huang and the proteomics core laboratory at Baylor College of Medicine for their assistance with the Luminex system. The illustrations in this paper were created using biorender.com.

