## Supplemental Data for "Shear Stress Induces a Time-Dependent Inflammatory Response in Human Monocyte-Derived Macrophages"

Supplemental Material

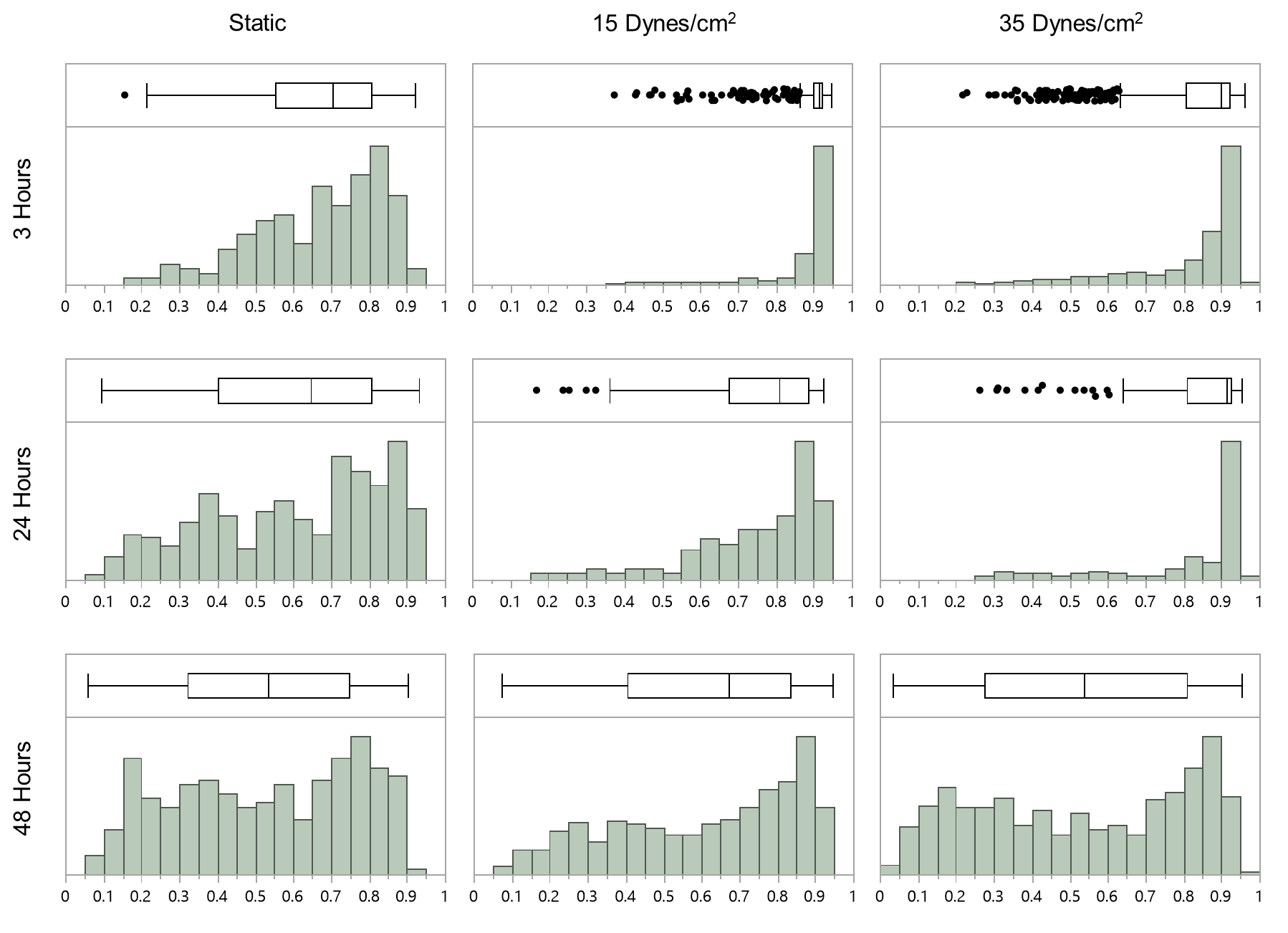

**Supplemental Figure 1.** Histograms and boxplots representing the circularity distribution of MDMs cultured under static, 15 dynes/cm^2^, and 35 dynes/cm^2^ conditions for a period of 3, 24, and 48 hours. The boxplots show the outliers, minimum, median, and maximum circularity for each group.

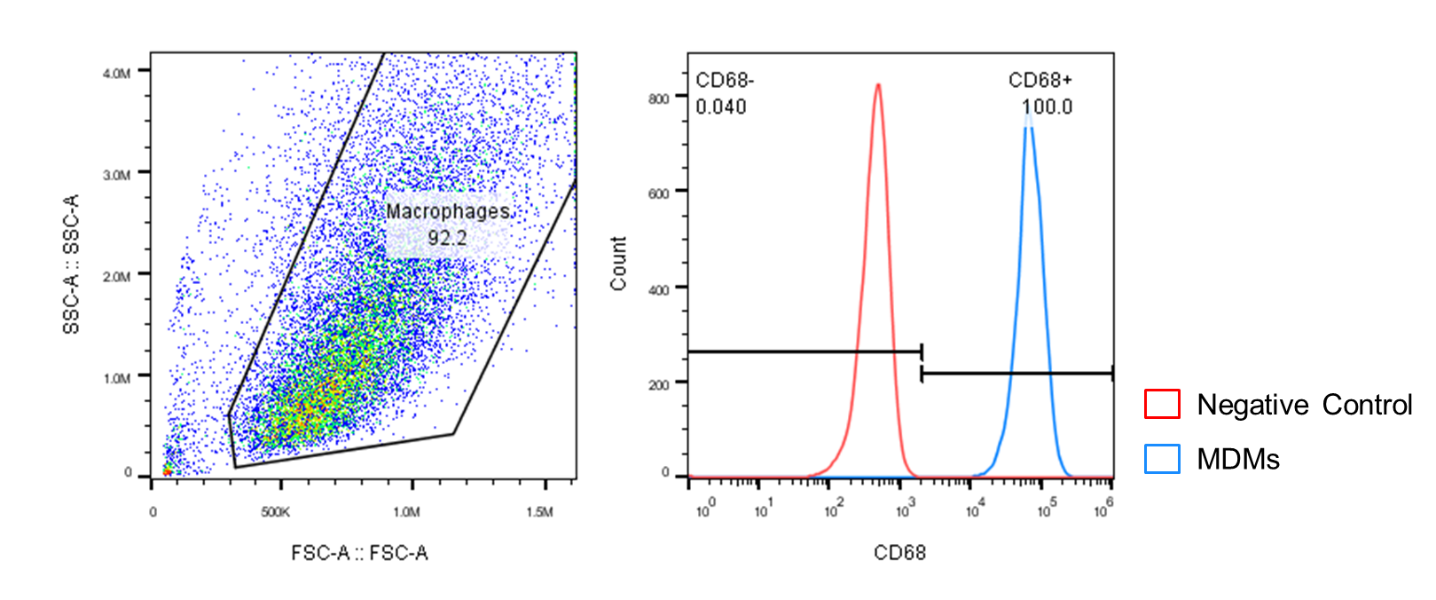

**Supplemental Figure 2.** Analysis of purity through flow cytometry. Demonstrates that 92% of the cells are positively expressing macrophage marker CD68.

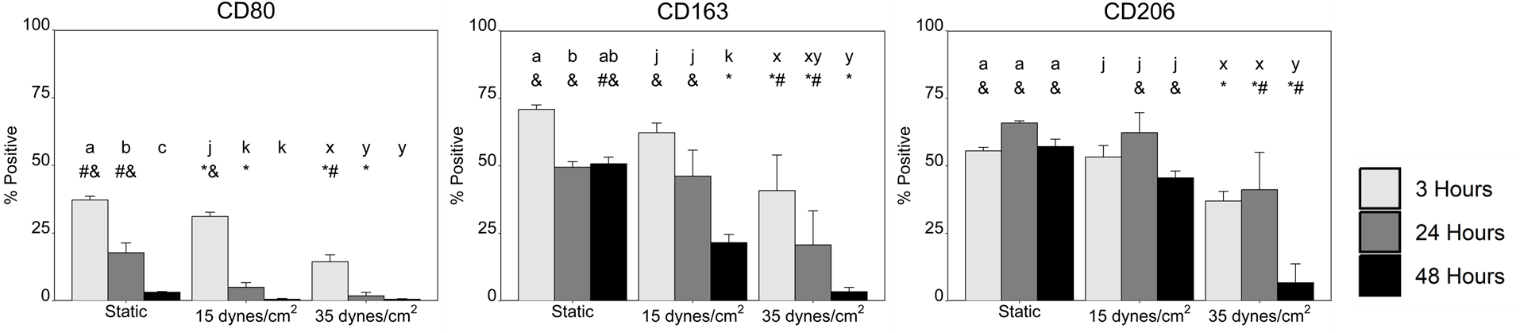

**Supplemental Figure 3.** MDM protein expression following shear exposure. Flow cytometry was performed to quantify percent positive protein expression of CD80, CD163, and CD206 following shear exposure. Data is represented as mean ± SD, n ≥ 3. Different letters indicate statistically significant differences between timepoints (p < 0.05) within a shear condition. Letter groups are as follows: a, b, c compares within the static condition; j, k, l compares within the 15 dynes/cm­^2^ condition; x, y, z compares within the 35 dynes/cm^2^ condition. Statistically significant differences between shear conditions within a timepoint are indicated by: *different from the static condition at the same timepoint; ^#^different from the 15 dynes/cm­^2^ condition at the same timepoint; ^&^different the 35 dynes/cm^2^ condition at the same timepoint.

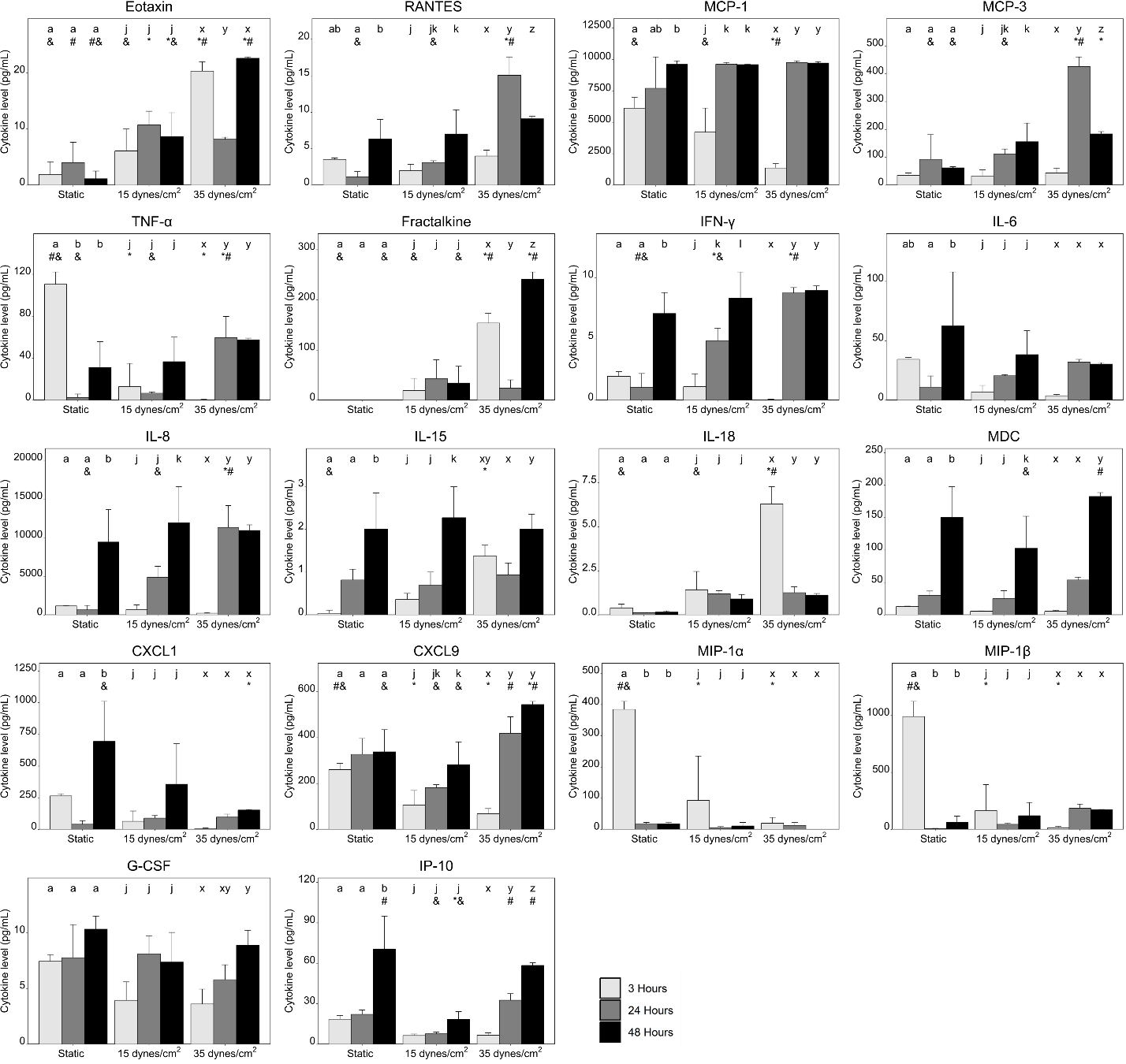

**Supplemental Figure 4.** Chemokines and pro-inflammatory cytokine secretion from MDMs cultured under static, 15 dynes/cm^2^, and 35 dynes/cm^2^ conditions for a period of 3, 24, and 48 hours. Data is represented as mean ± SD, n = 4. Different letters indicate statistically significant differences between timepoints (p < 0.05) within a shear condition. Letter groups are as follows: a, b, c compares within the static condition; j, k, l compares within the 15 dynes/cm­^2^ condition; x, y, z compares within the 35 dynes/cm^2^ condition. Statistically significant differences between shear conditions within a timepoint are indicated by: *different from the static condition at the same timepoint; ^#^different from the 15 dynes/cm­^2^ condition at the same timepoint; ^&^different from the 35 dynes/cm^2^ condition at the same timepoint.

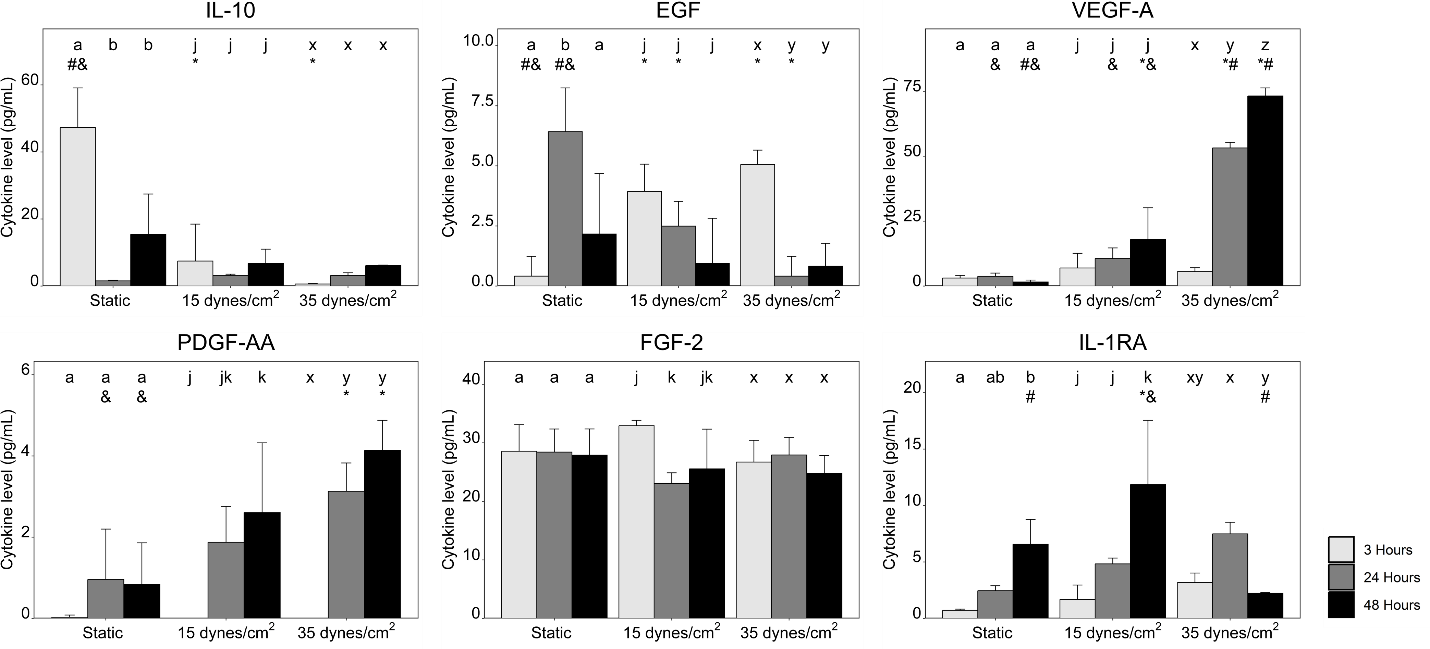

**Supplemental Figure 5.** Anti-inflammatory cytokine secretion from MDMs cultured under static, 15 dynes/cm^2^, and 35 dynes/cm^2^ conditions for a period of 3, 24, and 48 hours. Data is expressed as mean ± SD, n = 4. Different letters indicate statistically significant differences between timepoints (p < 0.05) within a shear condition. Letter groups are as follows: a, b, c compares within the static condition; j, k, l compares within the 15 dynes/cm­^2^ condition; x, y, z compares within the 35 dynes/cm^2^ condition. Statistically significant differences between shear conditions within a timepoint are indicated by: *different from the static condition at the same timepoint; ^#^different from the 15 dynes/cm­^2^ condition at the same timepoint; ^&^different from the 35 dynes/cm^2^ condition at the same timepoint.
